# When the host’s away, the pathogen will play: the protective role of the skin microbiome during hibernation

**DOI:** 10.1101/2023.11.28.568562

**Authors:** TS Troitsky, VN Laine, TM Lilley

## Abstract

The skin of animals is enveloped by a symbiotic microscopic ecosystem known as the microbiome. The host and microbiome exhibit a mutualistic relationship, collectively forming a single evolutionary unit sometimes referred to as a holobiont. Although the holobiome theory highlights the importance of the microbiome, little is known about how the skin microbiome contributes to protecting the host. Existing studies focus on humans or captive animals, but research in wild animals is in its infancy. Specifically, the protective role of the skin microbiome in hibernating animals remains almost entirely overlooked. This is surprising, considering the massive population declines in hibernating North American bats caused by the fungal pathogen *Pseudogymnoascus destructans*, which causes white-nose syndrome. Hibernation offers a unique setting in which to study the function of the microbiome because, during torpor, the host’s immune system becomes suppressed, making it susceptible to infection. We conducted a systematic review of peer-reviewed literature on the protective role of the skin microbiome in non-human animals. We selected 230 publications that mentioned pathogen inhibition by microbes residing on the skin of the host animal. We found that the majority of studies were conducted in North America and focused on the bacterial microbiome of amphibians infected by the chytrid fungus. Despite mentioning pathogen inhibition by the skin microbiome, only 30,4 % of studies experimentally tested the actual antimicrobial activity of symbionts. Additionally, only 7,8 % of all publications studied defensive cutaneous symbionts during hibernation. With this review, we want to highlight the knowledge gap surrounding skin microbiome research in hibernating animals. For instance, research looking to mitigate the effects of white-nose syndrome in bats should focus on the antifungal microbiome of Palearctic bats, as they survive exposure to the *Pseudogymnoascus destructans* -pathogen during hibernation. We also recommend future studies prioritize lesser-known microbial symbionts, such as fungi, and investigate the effects of a combination of anti-pathogen microbes, as both areas of research show promise as probiotic treatments. By incorporating the protective skin microbiome into disease mitigation strategies, conservation efforts can be made more effective.

## INTRODUCTION

Animals are constantly under attack from a plethora of microorganisms that have the potential to cause disease and even mortality. However, to infect an animal, these microbes have to first permeate the skin, which is the primary barrier between the host and the environment [1, 2]. The skin is a cool, acidic environment that is covered by sebaceous glands that secrete an antimicrobial substance, sebum. Sebum lubricates the skin and facilitates the growth of commensal microbes such as archaea, bacteria, viruses, and fungi. Together these microbes form a mutualistic community referred to as the skin microbiome [1, 3]. The host provides the symbionts with a favorable environment to propagate in and the microbes contribute by helping the host heal wounds, educating the immune system and preventing colonization of new microbes, which may have pathogenic properties [1, 2, 4, 5].

Microbiome research has increased in popularity in recent years with studies focusing mainly on the beneficial role of the gut, oral and skin microbiome of organisms, notably in humans [6, 7]. These studies give support to the holobiome theory, which suggests the host and its microbiome can be viewed together as a single evolutionary unit instead of separate entities [8, 9]. This perspective changes the definition of an individual to include the microorganisms living in and on the host. In many regards, the host cannot survive without its microbial symbionts, which also outnumber the cells of the host [10, 11]. This obligatory symbiosis also exists in animals, plants, and various other organisms [8]. Since the genomes of the microbes contributing to the microbiome evolve faster than the genome of the host, it can play a fundamental role in the host’s ability to rapidly adapt to environmental disturbances and new potentially pathogenic microbes [12]. This may be an important adaptation as climate change exposes species to novel pathogens.

The skin microbiome in particular is very sensitive to changes both in the environment and the host [1], which affects the holobiont’s ability to respond to changes. *Dysbiosis* or disruption in the composition of the skin microbiome can cause an imbalance that has a negative effect on host survival [2]. Dysbiosis often occurs when the amount of commensal microbes is reduced due to factors like immune deficiencies or exposure to pathogens, resulting in the microbiome losing its ability to protect the host [13]. For example, the diversity of the sheep (*Ovis aries*) skin microbiome is known to decrease preceding the onset of foot rot [14]. In addition, the artificial reduction of skin microbiome richness in salamanders before exposure to the deadly fungal pathogen that causes chytridiomycosis (*Batrachochytrium dendrobatidis,* hereafter *Bd*) leads to higher mortality [15]. In general, tropical amphibian species threatened by chytridiomycosis have lower skin bacterial diversity than non-threatened species [16].

On the other hand, the enrichment of certain microbes can be beneficial to the holobiont. Antimicrobial bacteria that inhibit pathogen growth *in vitro* have been found on the skin of amphibians, reptiles, fish, and mammals [17–20]. These bacteria, along with other protective microbes, are often referred to as probiotics. Testing the inhibition ability of these bacteria is becoming exceedingly popular in amphibians [21–23], because chytridiomycosis has caused major population declines in both the Americas and Eastern Australia [24, 25]. Mutualistic bacteria living on amphibian skin are known to produce antimicrobial agents, such as violacein and prodigiosin, that can inhibit the growth of *Bd* and suppress inflammation [13, 26, 27]. Thus, both positive and negative changes in skin microbiome composition seem to have a direct effect on the fitness of the host organism.

Due to its warm and moist nature, the skin provides an ideal environment for fungi to grow on, simultaneously making the skin more susceptible to fungal infections [28]. Over the past three decades, wildlife populations have experienced unprecedented, high-profile declines due to emerging infectious fungal diseases such as chytridiomycosis [29]. Another example of a deadly, skin-infecting mycosis that could potentially be treated with probiotics is white-nose syndrome (WNS) in insectivorous, hibernatory bats. WNS is caused by the psychrophilic fungus *Pseudogymnoascus destructans* (hereafter *Pd*), which invades and infects the skin causing a distinct fungal growth on the wings and muzzle of hibernating bats during winter [30, 31]. The fungal propagation arouses bats from torpor depleting their fat reserves, and eventually leading to starvation during a period when minimal insect-food is available. The disease was first discovered in the winter of 2006-2007 in New York and it has devastated Nearctic bat populations ever since, endangering once abundant species, such as the little brown bat (*Myotis lucifugus)* [32, 33].

The reason WNS has had such a calamitous effect on Nearctic bat populations can be attributed to the pathogen infecting bats when they are most vulnerable, during hibernation. The body temperature of hibernating bats drops drastically to resemble that of the ambient temperature in the hibernacula (2-14 °C) [30, 34]. Bats are, therefore, heterothermic, meaning they switch between an endothermic active state to an exothermic torpor state [35]. This radical change in thermoregulation is comparable to the ectothermic strategy of amphibians since both bats and their skin microbiome must tolerate substantial temperature fluctuations. This poses an added burden to both the host and its skin microbiome.

In addition, the metabolism and immune system of a bat become suppressed during hibernation, because they are energetically costly [36, 37]. This is exemplified by a significant decrease in the number of circulating leukocytes in the bloodstream during torpor [37]. Hibernation is an optimal strategy for insectivorous bats to save energy when food is scarce, and bats can remain torpid from days to months without eating [35]. However, the ability of the bat to defend itself against pathogens during this time becomes reduced due to its down-regulated immune system. Although most microscopic pathogens do not propagate well in cold temperatures [37], *Pd* thrives in the approximate temperature bats hibernate in, posing a significant threat [34].

However, not all bats get infected when exposed to *Pd*. In the Palearctic, where the fungus originates, bats tolerate exposure to the pathogen without infection or mortality [38, 39]. Species such as the greater mouse-eared bat (*Myotis myotis*) can tolerate high pathogen loads without apparent negative consequences [39, 40], suggesting the parasitic relationship has evolved into something that more resembles commensalism [41, 42]. One hypothesis to explain this phenomenon is that Palearctic bats have evolved a tolerance due to their longer history of exposure to the pathogen [43]. Molecular evidence implies Palearctic bats have been exposed to *Pd* for an extensive period of time, while Nearctic bats have had a mere 20-year bout with the pathogen since it was introduced from Europe [44, 45]. The protective skin microbiome could have enabled bat populations in the Palearctic to endure *Pd* exposure until the host develops tolerance.

Hibernation offers a unique setting in which to study the protective role of the skin microbiome because as the bat is in a torpid state, the microbiome may remain active. The symbiotic bacteria living on the skin of bats benefit from host survival, thus, it is not surprising that several of these bacterial strains have been found to have antifungal properties that may inhibit the growth of *Pd* [19, 46, 47]. For example, the bacterial genus *Pseudomonas* that is commonly found on bat skin has been shown to inhibit the growth of *Pd* both *in vitro* [46–49] and *in vivo* [50, 51]. Viewing Palearctic bats as holobionts that have coevolved together with *Pd* can help explain how selection might have favored bats harboring these antifungal bacteria in abundance on their skin. It is also noteworthy to mention that many other animals, such as some frogs, snakes, bears, rodents, birds, and fish possess the ability to hibernate, exposing them to similar risks as bats [52–57]. Therefore, studying the composition and antifungal potential of the skin microbiome during hibernation is an exclusive opportunity to better understand disease dynamics and the protective role of the skin microbiome in animals.

The aims of this review are to determine: *(i)* whether the protective skin microbiome of hibernating animals has been studied; *(ii)* whether experimental research studying pathogen inhibition of the skin microbiome has increased in the past years; and *(iii)* which antifungal microbes have been identified and studied the most? We emphasize the importance of experimental research because without inhibition assays and probiotic trials, the protective capacity of the microbiome remains speculative at best. To address these questions, we conducted a systematic review encompassing a range of publications examining the protective function of the skin microbiome in animals (Fig 1).

**Figure 1.**
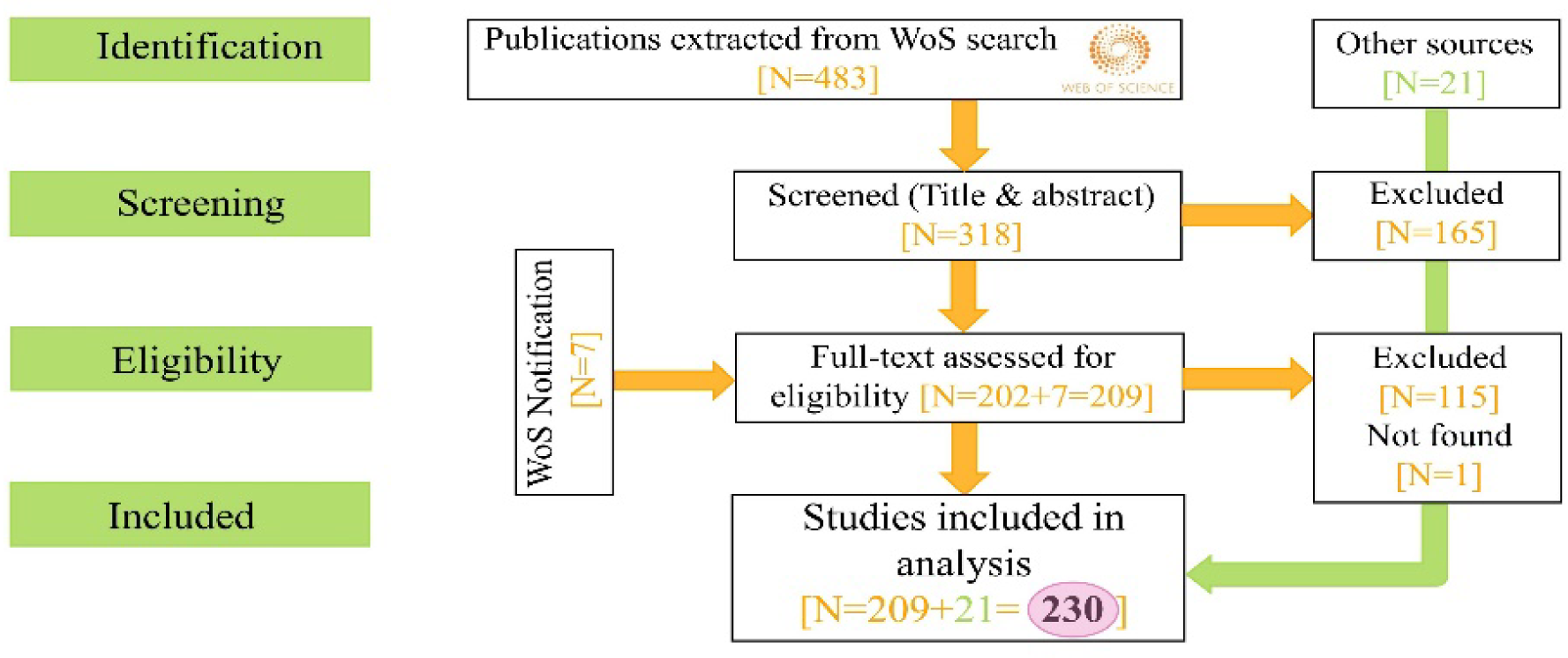
PRISMA diagram [99] explaining screening process of publications.

## MATERIALS AND METHODS

### 1) Systematic literature search

We performed a comprehensive keyword search on the *Web of Science* on the 13th of June 2023. Before defining the final search terms, we did multiple exploratory trials using different search words to determine which string of words would maximize the number of relevant references without adding an excessive number of irrelevant ones. For example, adding the words “probiotic” and “bioaugmentation” as a separate and obligatory search clause captured only 81 publications. Therefore, we added the words to the previous clause, making them facultative. We conducted the final optimized search using the following terms:

ALL=(“microbiota” OR “microbiome”) AND ALL=(“skin” OR “cutaneous” OR “epidermis” OR “dermal”) AND ALL=(“resistance” OR “inhibit” OR “antifungal” OR “pathogen” OR “fungal” OR “bioaugmentation” OR “probiotic”) AND ALL=(“vertebrate” OR “invertebrate” OR “animal” OR “mammal” OR “reptile” OR “amphibian” OR “fish” OR “bird” OR “bat”)

This yielded 483 publications ranging from the years 2005-2023 that were screened by TST according to the PRISMA diagram (Fig 1.). Articles were found suitable for this review based on the following inclusion criteria: (i) they studied the skin microbiome (as opposed to just gut or oral microbiome); (ii) they mentioned skin-infecting pathogens and antimicrobial symbionts living on the skin of the host; (iii) they studied non-human animals. Reviews and publications that did not meet these criteria were excluded, including one publication that was not accessible. In addition to this, we added 21 publications found elsewhere that fit the search criteria and seven publications that we were notified about by *Web of Science* alert, resulting in the final data set (N=230).

### 2) Metadata extraction

We extracted metadata from all relevant references for the final database. We documented the geographical and taxonomic range of the studies, the host’s captivity status, whether the study solely examined microbiome composition (descriptive) or also assessed the microbiome’s response to pathogens (experimental), as well as how the microbiome’s response was tested and whether pathogens were known to infect hosts during hibernation. This was done by cross referencing literature and/or checking pathogen propagation temperatures (if optimal pathogen propagation temperature was not similar to the temperature in hibernacula, the pathogen was not considered a threat during hibernation). For studies that experimentally tested microbes against pathogens we also determined the type of pathogen and antimicrobial genera detected on skin and whether the microbes were successful in inhibiting the pathogens.

### 3) Data visualization and statistical analyses

We performed data analysis and visualizations using R version 4.2.2 [58] using packages ‘ggplot2’ version 3.4.2 [59] and ‘bipartite’ version 2.18 [60]. Additionally, we used Inkscape version 1.3 [61] to edit the visualizations. We used a binomial generalized linear model to analyze how many of the publications actually experimentally tested the pathogen inhibition ability of the microbiome, in proportion to all published studies over the past years (glm(formula = cbind(experimental, descriptive) ∼ year, family = “binomial”)). We excluded the year 2023 from the analysis, since the year is not over, and more studies are likely to be published before the end of the year.

## RESULTS

### General summary of literature

In our initial *Web of Science* search, we identified 483 publications. Following the screening of titles and abstracts, 318 were considered relevant and underwent full text inspection. Among these, 202 met our inclusion criteria and seven publications were added after a Web of Science alert. An additional 21 studies were added from other sources, resulting in 230 publications (Supplementary file 1).

The majority of skin microbiome studies were conducted in the Western Hemisphere, with 59,1 % of studies taking place in North America and 8,3 % in South America (Fig 2). The remaining 32,6 % of studies were spread between Europe (13,5 %), Asia (13,0 %), Africa (2,2 %), and Oceania (3,0 %). Additionally, two studies (0,9 %) sampled animals from multiple continents.

**Figure 2.**
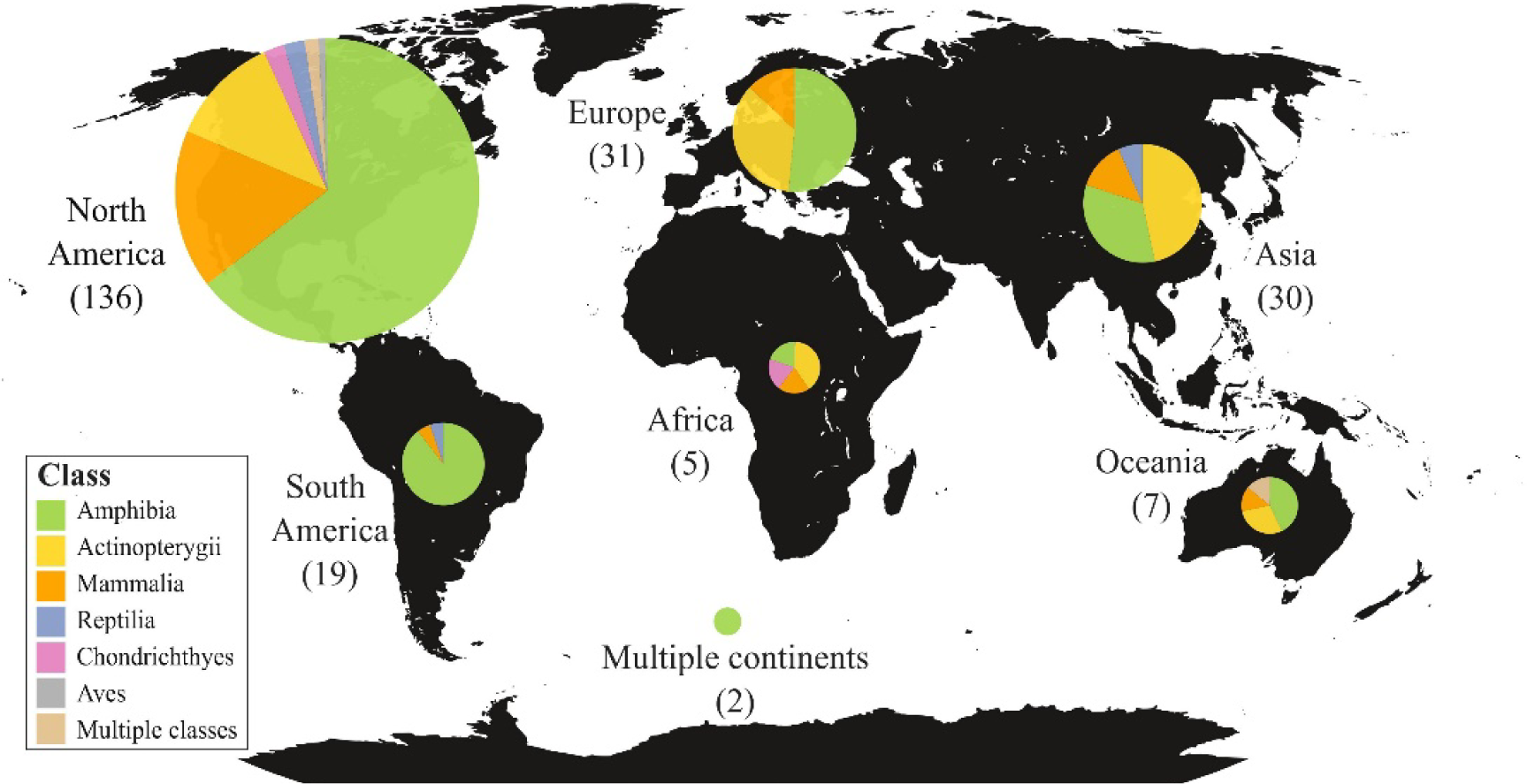
Summary of literature. The proportion of skin microbiome studies conducted on different animal classes on all continents. Number of studies in parentheses.

Predictably, the most studied animal class was amphibians (59,6 %), followed by ray-finned fishes (19,6 %), and mammals (14,8 %). The remaining 6,0 % of studied species were divided into reptiles (2,6 %), cartilaginous fishes (1,7 %), and birds (0,4 %) with three studies investigating multiple classes (1,3 %). About half (52,0 %) of the publications studied the skin microbiome of wild animals, while 42,3 % focused on captive animals, and 5,7% considered both.

Despite the search set to contain at least one word regarding pathogen inhibition (“antifungal”, “pathogen”, “resistance”, “inhibit”, “fungal”, “bioaugmentation” or “probiotic”), most of the publications (69,6 %) only described the skin microbiome composition of the host, without testing the inhibition ability of potentially antifungal bacteria found on the skin. Altogether, only 30,4 % of publications tested inhibition ability by either conducting inhibition assays *in vitro* or testing probiotic treatments *in vivo*. Out of these studies 65,2 % tested inhibitory ability using inhibition assays, 24,2 % used probiotic treatment, and 10,6 % used both. Among these studies 60,6 % experimented on antifungal amphibian symbionts, 21,2 % on mammalian symbionts, 16,7 % on fish symbionts, and 1,5 % on reptile symbionts. The majority (54,7 %) of experimentally studied host species were captive, while 37,5 % of publications studied wild animals, and 7,8 % studied both.

#### i) Has the protective skin microbiome of hibernating animals been studied?

Only 18 publications (7,8 % of all articles) studied the protective microbiome during hibernation. Six of these studies were solely descriptive and 12 were experimental. Although many amphibian and reptilian species are capable of hibernation, no studies were conducted on the protective role of their skin microbiome during hibernation. In fact, all studies that sampled the skin microbiomes of hibernating animals involved bats and *Pd*.

Among the 12 experimental publications that focused specifically on a pathogen that infects hosts during hibernation, nine studies tested antifungal microbes against *Pd* using inhibition assays *in vitro* and three studies tested probiotic treatments *in vivo*. Only two of these studies were conducted in the Palearctic (Germany and China), where bats survive exposure to *Pd* without infection [49, 62]. Both studies used bacteria in inhibition assays to successfully suppress the growth of *Pd.* The remaining ten studies were conducted in North America.

Altogether, only seven studies have been published about the skin microbiome of Palearctic bats. Among these, five were descriptive [63–67] and two experimental [49, 62]. Four took place in China [49, 63, 64, 66], while the remaining three were conducted in Germany [62], Poland/Armenia [65], and Belgium [67]. Out of these, only five publications sampled hibernating bats [49, 63–66], while one study sampled active bats and the wall of the hibernacula [67], and one acquired the symbiont tested against *Pd* from the environment (not bat skin) [62].

#### ii) Has experimental skin microbiome research studying pathogen inhibition increased in recent years?

The overall amount of research on the protective role of the skin microbiome in non-human vertebrates has increased dramatically over the past 18 years, with the first study conducted in 2005 (Fig 3). Regardless of the growing interest in this field, the proportion of experimental studies investigating inhibitory ability of antimicrobial microbes residing on the skin has decreased significantly (p < 0,001, −0.20187 ± 0.04862) in proportion to the number of studies published. This might be explained by the fact that the topic is vastly unexplored, and most studies focus on solely describing the skin microbiome composition of animals and whether known antimicrobial taxa are found on the skin. It is, however, noteworthy to mention that the number of all studies published (both descriptive and experimental) has been lower in the year 2023 compared to previous years. It remains to be determined whether more studies will be published by the end of the year.

**Figure 3.**
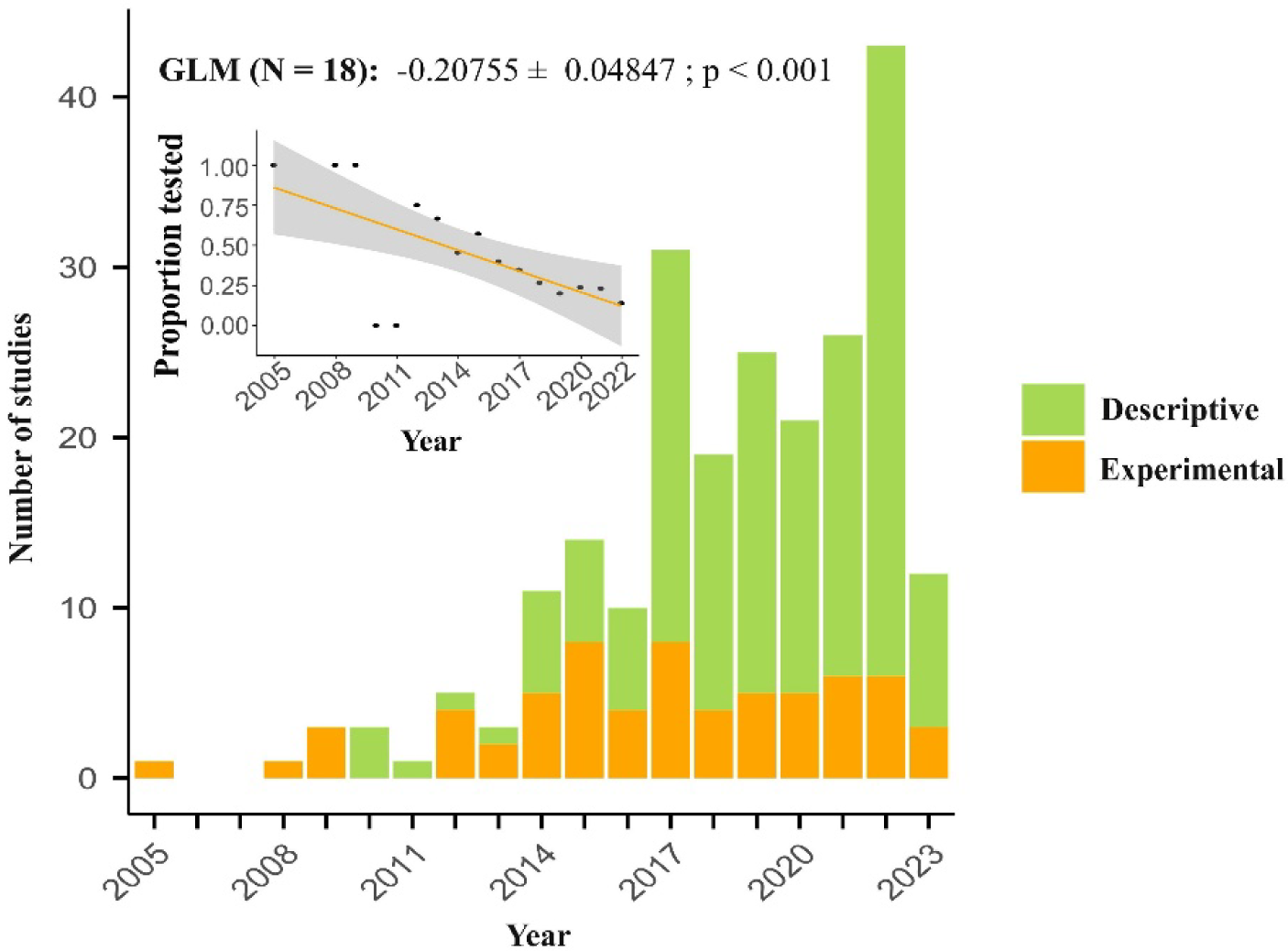
Temporal trends in protective skin microbiome studies in non-human animals. The year 2023 was excluded from the GLM and scatter plot since the year is not over.

#### iii) Which antifungal microbes have been identified and studied the most?

A total of 105 microbial genera were found to show weak to strong pathogen inhibition in the experimental studies. The majority of tested microbes were bacteria (84,8 %), but fungi (13,3 %), and archaea (1,9 %) were also tested successfully. Out of these, 51 genera were experimentally tested more than once, and 27 genera were tested on two or more classes of animals (Fig. 4).

**Figure 4.**
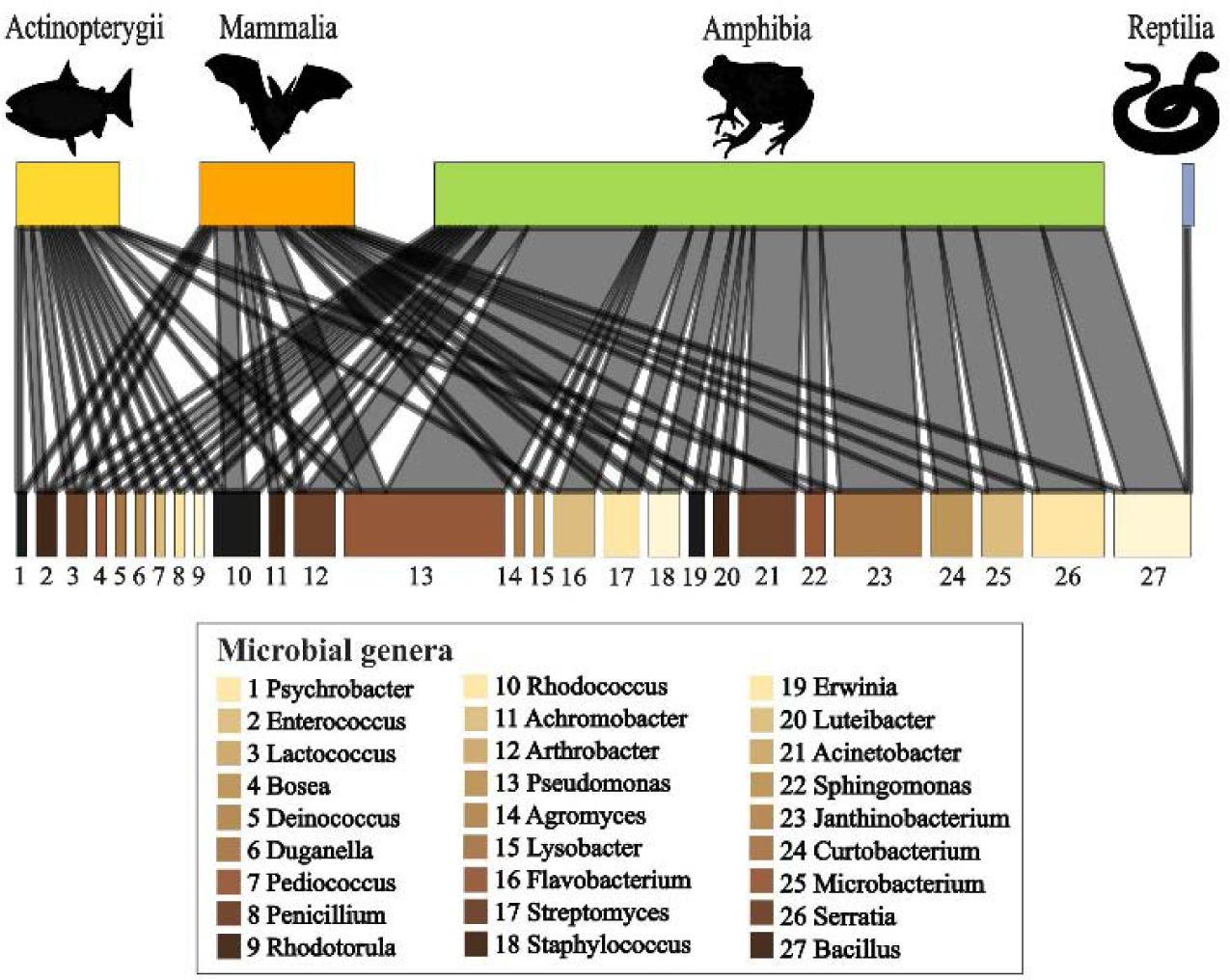
Antifungal microbes found on the skin of two or more classes of animals that successfully inhibited pathogens. Size of block indicates number of studies conducted (larger block = more studies).

The most popular bacterial genus tested against pathogens was *Pseudomonas*. It was tested against multiple pathogens (*Pd, Bd,* and others) that infect several classes of animals including mammals, amphibians, and ray-finned fishes. *Pseudomonas* species showed strong to moderate inhibition of pathogens and they are considered one of the primary candidates for *Pd*-inhibition in bats [46, 68]. Altogether, 31 studies testing *Pseudomonas* showed successful pathogen inhibition, however, two studies using *Pseudomonas* as a probiotic were unsuccessful [22, 69]. The authors of one failed trial noted that the skin microbiome still retained a defensive role against *Bd*, but that the antifungal isolates were unable to colonize the skin of the amphibian host. This issue may be addressed in future experiments by reapplying treatment or prolonging exposure to treatment [22].

Other microbes successfully tested against pathogens in multiple studies included the genera *Janthinobacterium, Bacillus, Chryseobacterium, Stenotrophomonas, Serratia, Acinetobacter, Rhodococcus,* and *Enterobacter*, indicating these bacteria show promise as probiotics and should be investigated in more detail. However, it is important to mention that *Janthinobacterium*, *Chryseobacterium*, *Stenotrophomonas,* and *Rhodococcus* also failed some trials. For example, while the genus *Rhodococcus* showed strong inhibition of *Pd in vitro* [70], a recent *in vivo* experiment on bat skin was not successful [71]. These results highlight the need for more trials to determine whether these microbes can, in fact, be used in wildlife disease mitigation. Further experiments will also help assess the microbial mechanisms of inhibition, which provide important information about the conditions that best facilitate pathogen inhibition.

Not all experimental publications studied the inhibitory effects of a single microbial strain, six publications (9,1 % of experimental studies) researched the collective effect of a group of inhibiting microbes. Bacterial genera used in the consortium studies were *Pseudomonas, Janthinobacterium, Bacillus, Chryseobacterium, Stenotrophomonas, Serratia, Acinetobacter, Enterobacter, Microbacterium, Staphylococcus, Citrobacter, Comamonas, Pedobacter, Chitinophaga, Iodobacter*, *Collimonas, Curvibacter,* and *Sanguibacter.* Out of these, five publications were successful in inhibiting pathogens [20, 72–75], and one was not [69].

## DISCUSSION

In this review, we illustrate that the protective role of the skin microbiome is becoming an increasingly popular topic of research. However, most publications focus on describing the composition of the skin microbiome and identifying known antifungal microbes without testing their pathogen inhibition ability. This phenomenon may be explained by the novelty of the topic, as the majority of publications aim to simply describe the microbial diversity on the skin of hosts, before experimentally testing them against pathogens. Additionally, the knowledge attained so far centers mainly around amphibians in the Nearctic, leaving other animal classes and continents overwhelmingly unexplored. The popularity of protective skin microbiome studies in North American amphibians can be explained by the disastrous emergence of chytridiomycosis in 1993 [24] and the uneven distribution of research funding opportunities that are overly represented in the Nearctic [76].

The main research gap we want to highlight with this review is the lack of publications about the protective role of the skin microbiome in hibernating animals, specifically studies that experimentally test the inhibition ability of cutaneous microbiota against pathogens. The suppression of the host’s immune system during hibernation amplifies the importance of the skin microbiome since it may remain active when the host is not. It would be especially beneficial to study the protective role of the skin microbiome in species of animals that survive exposure to pathogens during hibernation, such as Palearctic bats. So far, the skin microbiomes of only 13 of over 100 bat species in the Palearctic have been studied. Most of these studies focused on fungal symbionts and just one study was conducted on the protective mycobiome of *M. myotis,* the flagship species known to tolerate high *Pd* loads in Europe. To our knowledge, there are no published data on the mutualistic bacteria living on the epidermis of *M. myotis*. In fact, the bacterial composition of most Palearctic bat species and possible temporal changes in their microbiome composition (for example during hibernation) remain unknown.

### Are probiotics the solution to lethal skin disease in wildlife?

Describing and experimentally testing microbial species found on host skin are the first steps to developing a non-toxic disease mitigation strategy for lethal skin infections in wild animals. As mentioned earlier, using antimicrobial bacteria as a preventative probiotic treatment on the skin to help mitigate disease has already been explored in some organisms [26, 50, 77]. Results from these studies have varied, however, most studies have found encouraging findings in several classes of animals. For example, the bioaugmentation of a known antifungal bacterium (*Janthinobacterium lividum)* on frog skin successfully prevented mortality due to chytridiomycosis [17]. Moreover, probiotic treatments tested on walleye fish (*Sander vitreus*) were found to have a significant antagonistic effect against a common pathogen (*Flavobacterium columnare*) and increase the survival of fish exposed to the pathogen [78]. In addition, a probiotic bacterium isolated from feline skin successfully reduced the colonization of a pathogen when added to the epidermis of mice, indicating certain probiotics could be effective across multiple species [79]. While this provides compelling evidence for the justification of probiotic use, other publications have reported contradicting results [22, 69, 80], suggesting more information is needed before probiotic treatments can be successfully applied to wildlife disease mitigation.

Our results indicate that several microbial species, mostly bacteria, have been shown to exhibit potential as probiotics. Notably, the bacterial genus *Pseudomonas* has demonstrated the inhibition ability of several pathogens infecting multiple classes of animals, including bats and *Pd* [46, 81–83]. However, there are various other microbial genera that have shown inhibition ability but are still overlooked. Fungi are among the often disregarded species that have also shown promise in pathogen inhibition [84–86]. For example, North American bat species resistant to WNS exhibit a more diverse cutaneous mycobiome compared to WNS-susceptible species [86]. Some common fungal genera identified on bat skin, such as *Cutaneotrichosporon*, *Aureobasidium*, and *Holtermanniella*, have also been found to inhibit the growth of *Pd in vitro*, albeit weakly [86, 87]. Additionally, gram-positive bacteria may be overlooked in these studies since DNA extraction methods do not always successfully permeate the thick outer layer of the bacteria [88, 89]. These bacteria may also possess the ability to inhibit pathogens, but could be underrepresented in these datasets and, therefore, not tested for inhibition.

It is also important to acknowledge that certain mutualistic microbial genera, such as *Pseudomonas*, are known pathogens for certain organisms [20, 90, 91], meaning the effect of the microbial genus is highly dependent on context [21]. For example, *Pseudomonas fluorescens*, a commensal on bat skin [51], can be lethal to fruit flies (*Drosophila melanogaster*) and ladybird beetles (*Henosepilachna vigintioctopunctata*) [92]. Timing of treatment is also of importance since the addition of *P. fluorescens* to bat skin before exposure to *Pd* increased disease severity, while simultaneous treatment and exposure reduced *Pd* invasion [50]. When utilizing probiotics, there is always the risk that the symbionts could spread to and infect non-target species causing more harm than good. Hence, it is advisable that the probiotic is indigenous to the local environment and has been studied adequately before adding treatment to an ecosystem or species [21, 93].

In addition to testing the pathogen inhibiting ability of just one microbial strain, there seems to be an emerging trend of testing a consortium of bacteria against pathogens. Multiple studies have found that more diverse communities of bacteria can outperform single strains in inhibiting pathogen growth [73–75]. Bacterial growth rate *in vitro* has also been found to be higher, when bacterial strains were grown together, instead of individually [72]. This is understandable given that a diverse community of organisms is known to be more resistant to invasions on both a macro-[94] and micro-scale [95]. For instance, as an analogous example, grassland plots with higher species diversity are more resistant to colonization by invasive plants than homogenous plots [94]. The interactions of microbial species within the microbiome mirror those of organisms in a macro-level ecosystem (for example a forest), which is why diversity means better pathogen resistance in the skin microbiome as well [1, 86].

### Future threats and conservation

As climate change progresses and humans encroach further into wildlife habitats, people and wildlife alike will be more regularly exposed to new potential pathogens [96, 97]. Fungi, in particular, should be treated with concern as fungal infections are notoriously difficult to treat due to their resilient nature. Over 600 species of fungi are known to infect vertebrates and many species have been identified as the causal agents of potential emerging infectious diseases (EIDs) in recent years [96, 98]. In fact, fungi are more closely related to animals than bacteria, and therefore, do not respond well to common antimicrobial treatments that work on bacterial infections [98]. WNS and chytridiomycosis have demonstrated how rapidly fungal disease outbreaks can devastate wildlife populations and highlight the need for preventative disease mitigation strategies.

It is often difficult to manage disease outbreaks in endangered wildlife populations, so captive breeding and reintroduction are occasionally used to attempt to restore declining populations [99]. These attempts are often costly and have varying success rates. In this review, the majority (54,7 %) of experimental skin microbiome studies were conducted on captive animals. However, since the skin microbiome is heavily influenced by the environment [1, 100], the results from these studies may not always be applicable to wild animals. For example, the skin microbiome of captive amphibians is known to be less diverse than that of their wild counterparts, which may become an issue when reintroducing captive animals back into the wild during conservation efforts [101]. Considering reduced diversity in the skin microbiome affects the host’s ability to resist infection, the holobiont perspective could be beneficial when planning and upgrading conservation methods [99].

## CONCLUSIONS

While the skin microbiome holds tremendous potential for disease mitigation, its protective role during hibernation is highly understudied. Not only is there a scarcity of publications describing the microbial diversity inhabiting the skin, but there is also a notable absence of experimental studies determining which microbes effectively inhibit pathogens. Hibernatory bats and WNS provide an exceptional study system for addressing this knowledge gap and we encourage researchers to tackle this subject by exploring the microbial species living on bat skin and their potential as probiotics in WNS mitigation. Specifically, the skin microbiome of Palearctic bats should be studied to determine how they survive exposure to *Pd*, as this information could be beneficial for solving the WNS crisis in North America. In particular, we recommend future research concentrate on testing the anti-pathogen activity of lesser-known symbionts, such as fungi, in addition to testing a consortium of known antifungal bacteria. We emphasize the importance of adopting a holistic approach which incorporates the holobiont perspective into conservation planning for more efficient results in disease mitigation.

## LIST OF ABBREVIATIONS

*Bd*: *Batrachochytrium dendrobatidis*
WNS: White-nose syndrome
*Pd*: *Pseudogymnoascus destructans*

## DECLARATIONS

### Ethical Approval

Not applicable.

### Authors’ contributions

TML, VNL and TST conceived the idea and designed the methodology. TST extracted data from the literature, analyzed the data, prepared the figures, and led the writing of the manuscript. All authors read, contributed to, and approved the final manuscript.

## Acknowledgements

The authors would like to thank Dr. Melissa Meierhofer for her insightful and constructive input in the selection of figures and models used in this review.

## Availability of data and materials

All data presented in the supplementary material.

**Supplementary file 1:** Data set (Review_Data_Troitsky.xlsx)

Contains list of chosen publications and data extracted from them that was used in model and figures.

